# Excitatory and inhibitory L2/3 neurons in mouse primary visual cortex are balanced in their input connectivity

**DOI:** 10.1101/2020.04.21.053504

**Authors:** Alexander P.Y. Brown, Lee Cossell, Troy W. Margrie

## Abstract

Quantitatively characterising brain-wide connectivity of neural circuits is of vital importance in understanding the function of the mammalian cortex. Here we have designed an analytical approach to examine data from hierarchical segmentation ontologies, and applied it in the comparison of long-range presynaptic connectivity onto excitatory and inhibitory neurons in layer 2/3 (L2/3) of mouse primary visual cortex (V1). We find that long-range connections onto these two general cell classes in L2/3 originate from highly similar brain regions, and in similar proportions, when compared to input to layer 6. These anatomical data suggest that distal information received by excitatory and inhibitory networks is highly homogenous in L2/3.

## Introduction

A quantitative characterization of inter- and intra-region connectivity is crucial in order to elucidate general principles and region-specific features of circuit structure and function. In the neocortex, information is processed by local networks of excitatory glutamatergic principal neurons and inhibitory GABAergic interneurons (Markram et al., 2004). The local circuit connectivity and response properties of excitatory and inhibitory neurons have been thoroughly studied in L2/3 of rodent V1 (Mason et al., 1991; Holmgren et al., 2003; Yoshimura & Callaway, 2005; Runyan et al., 2010; Bock et al., 2011; Ko et al., 2011; Hofer et al., 2011; Cossell et al., 2015; Wertz et al., 2015; Lee et al., 2016; Kim et al. 2018; Znamenskiy et al., 2018). For instance, although GABAergic inhibitory neurons make up only ∼20% of the cortical population, connections between excitatory and inhibitory neurons are much more dense (Hofer et al., 2011; Packer & Yuste, 2011) than connections between excitatory neurons (Holmgren et al., 2003). Moreover, local connectivity shows a high degree of specificity between functionally-related neurons (Bock et al., 2011; Ko et al., 2011; Cossell et al., 2015; Wertz et al., 2015; Lee et al., 2016; Znamenskiy et al., 2018). However, despite this knowledge of connectivity at a local scale, fewer studies have examined the specificity and pattern of brain-wide, long-range input onto specific cell types in V1, and the majority of those have looked at deeper layers 5 and 6 (Vélez-Fort et al., 2014; Kim et al., 2015; Liang et al. 2019).

Retrograde modified-rabies-virus (RV) tracing, which permits identification of monosynaptically coupled presynaptic cells, has been particularly useful for mapping long-range connectivity (Wickersham et al., 2007; Luo et al. 2018). Since the modified virus is intrinsically replication incompetent, presynaptic labelling is limited to neurons directly connected to the initial rabies-infected cells. Using Cre-driver lines for population tracing from molecularly defined cell types across different layers, previous RV tracing experiments have indicated V1 directly receives a variety of non-visual, multimodal sensory and spatial information (Vélez-Fort et al., 2014; Kim et al., 2015; Wertz et al., 2015; Leinweber et al., 2017; Liang et al., 2019). Further, single-cell and population tracing experiments from physiologically characterized neurons show that different functional classes of cells have partially non-overlapping input profiles (Vélez-Fort et al., 2014; Kim et al., 2015; Liang et al., 2019). Therefore, the anatomical origin of input onto deep layers of V1 is widespread and relates to the functional properties of postsynaptic neurons, but the extent to which these connectivity profiles differ according to specific target excitatory and inhibitory pathways in L2/3 is not known.

Given the differences in response properties, local circuit connectivity and molecular markers, a corresponding difference in the brain-wide input profile to excitatory and inhibitory cells in the cortex may also be expected. Therefore, here we have examined the extent to which long-range input profiles to glutamatergic and GABAergic populations in L2/3 differ. We developed an analytical approach, designed specifically for hierarchical segmentation ontologies, to directly compare these profiles in individual brains. Using this approach, we demonstrate that excitatory and inhibitory cells in L2/3 receive relatively similar long-range input, arising from the same brain regions and in similar proportions. Further, we show that the input maps onto different cell classes in L2/3 are quantitatively more similar than input maps to a class of principal cells in L6.

## Results

In order to characterise presynaptic input onto excitatory and inhibitory neurons in primary visual cortex (VISp), we used modified rabies-virus (RV) tracing (Wickersham et al., 2007). To specifically target initial transfection to GABAergic cells in L2/3 we injected Cre-dependent AAV superficially (∼120μm below the pia) in Gad2-Cre and Parv-Cre transgenic mice (referred to throughout as “gadOn” and “parv”, respectively) (**Fig. 1a**). To target L2/3 glutamatergic, excitatory cells, we instead used a Cre-Off AAV in Gad2-Cre mice (“gadOff”) (Saunders et al., 2012), so that the AAV was selectively expressed in Gad2-negative, putative excitatory neurons. Finally, we compared these datasets to a reanalysed dataset, which was generated by injecting the Cre-dependent AAV in Ntsr1-Cre transgenic mice (“ntsr1”) (Vélez-Fort et al., 2014), in which a subset of L6 excitatory cells are labelled.

**Figure 1.**
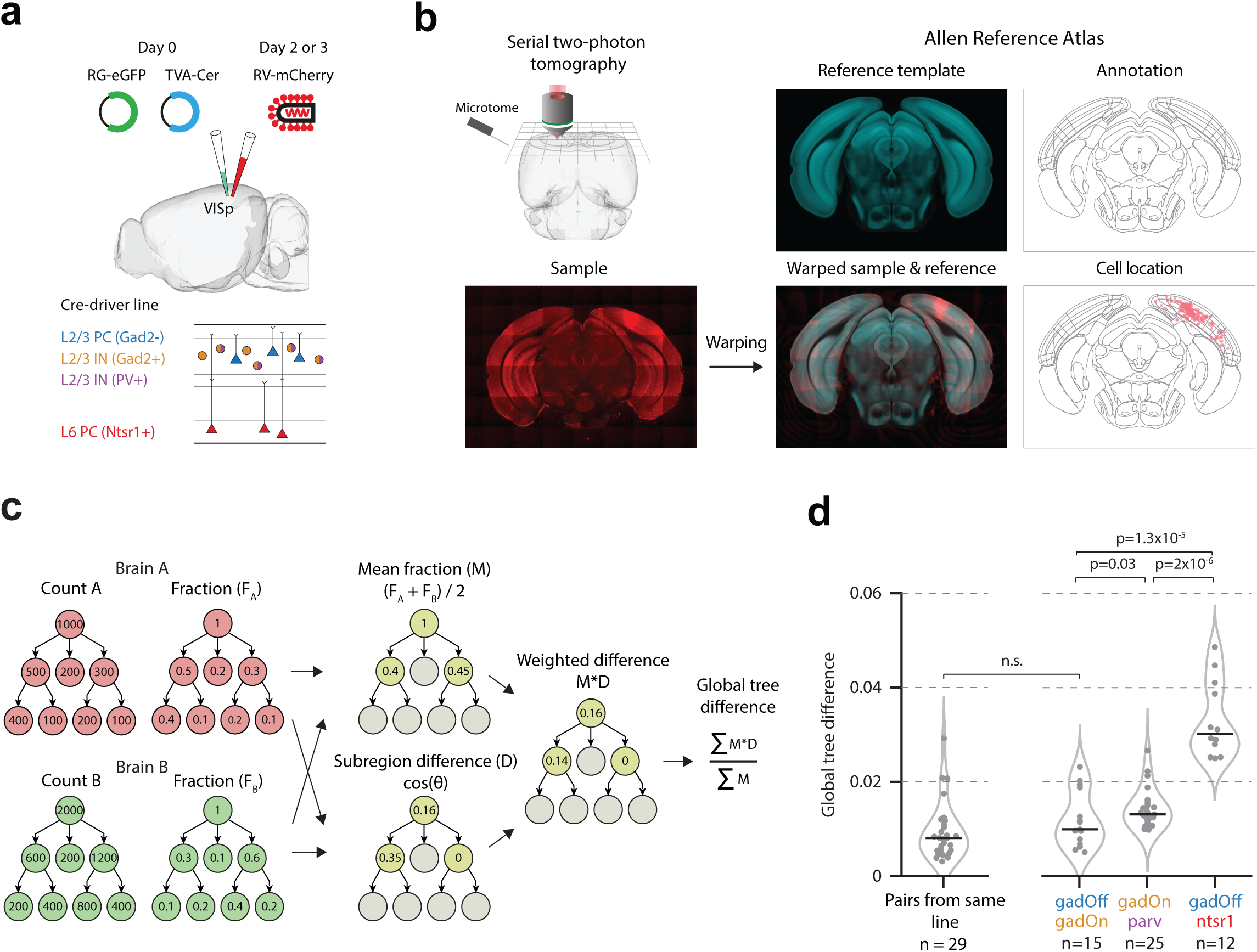
Global tree difference showing similarity of input to L2/3 relative to L6. (**a**) Schematic of experimental procedure. (**b**) Serial two-photon tomography was performed on each brain, and samples were aligned to and segmented according to the Allen Reference Atlas (Niedworok et al., 2016). (**c**) Outline of the algorithm used for calculating the GTD. For each brain, the cell counts in each region are converted to a fraction of the total labelled inputs. For a pair of brains, for each region, the mean fraction, M, is then calculated, as well as the cosine difference (D) between the distribution of cells within the region’s subregions. Regions with only a single subregion are excluded from further processing. Next, for each region, the mean fraction is multiplied by the cosine difference, to give a weighted difference score (M*D). Finally, this weighted difference score is summed across all regions in the hierarchy and normalized by the sum of the mean fractions across all regions. (**d**) Global tree difference for each pair of brains from within the same line (left), and for pairs of brains in gadOff and gadOn experiments, gadOn and parv experiments, and gadOff and ntsr1 experiments (right).

Following *ex vivo* imaging using serial two-photon tomography and cell counting, each brain was segmented, and presynaptic cells were assigned to brain regions using aMAP (Niedworok et al., 2016) with the Allen Reference Atlas ontology (**Fig. 1b**). In total, data were acquired from 17 brains (gadOn: n = 5; parv: n = 5; gadOff: n = 3 mice; ntsr1: n = 4), and the number of labelled presynaptic cells ranged widely across experiments (gadOn: 691-7217; parv: 993-16944; gadOff: range 420-2296; ntsr1: 1112-6266). As the majority of labelled cells were ipsilateral to the injection site in all lines (percentage ipsilateral inputs: gadOn: 97.1 ± 2.1%; parv: 99.3 ± 0.2%; gadOff: 99.8 ± 0.1%; ntsr1: 98.0 ± 0.4%; mean ± SEM; **Supplementary Fig. 1**), we restricted further analysis to the ipsilateral hemisphere.

We first sought to explore the differences in input connectivity onto excitatory and inhibitory cells in L2/3. The detailed analysis of brain-wide long-range connectivity data entails a number of distinct and largely unexplored analytical challenges. Firstly, the hierarchical nature of segmentation ontologies necessitates difficult and often arbitrary selection of regions for analysis. There may well be no clear criteria on which to decide at which level of granularity to analyse the data – whether to assign cells to the finest level of detail possible in the particular classification, or whether to agglomerate small subregions in to a single parent region. Secondly, the number of brain regions in modern ontologies makes addressing the problem of multiple comparisons critical in any detailed analysis of segmentations.

We therefore designed our analyses to address these concerns. First, in order to directly compare the distribution of cells amongst brain regions in two individual brains – such as presynaptic connectivity maps – we developed a pairwise scalar difference measure (the “global tree difference”; GTD) (**Fig. 1c**; **Supplementary Fig. 2**) which utilizes the entire hierarchy of an ontology. For each segmentation in a given pair, the fraction of total presynaptic input was calculated in each region of the hierarchy (**Fig. 1c**, left). Each brain region was then assigned a cosine distance score based on the difference in the distribution of cells amongst its child subregions in the two brains (the subregion difference, D) (**Fig. 1c**, middle). Next, an average fractional connectivity map was calculated for the two brains (the mean fraction, M) (**Fig. 1c**, middle), and used to calculate a weighted cosine distance. These calculations were summed and normalized to give a scalar measure of the difference between two individual brains (the GTD) (**Fig. 1c**, right).

If long-range connectivity is cell-type dependent, the profile of presynaptic labelling in experiments with the same population of starter cells should be more similar than experiment pairs with starter cell populations of different cell types. Thus, in order to establish a lower bound for the GTD, the value was first computed for all pairs of brains from within the same experimental condition (median GTD = 0.008, IQR = 0.006; n = 29 pairs; **Fig. 1d**, left). Next, in order to directly compare the pattern of input connectivity onto excitatory and inhibitory cells in L2/3, we computed the GTD for all pairs of brains in the gadOff and gadOn experiments (median GTD = 0.010, IQR = 0.010, n = 15) (**Fig. 1d**, right), and this value was not significantly different to the GTD value from pairs within the same experimental condition (p = 0.09; Wilcoxon rank-sum test). We next examined all pairs from the gadOn and parv experiments and found a significantly higher GTD value (median GTD = 0.013, IQR = 0.003, n = 25; p = 0.03), suggesting that presynaptic connectivity maps from the gadOff and gadOn lines are more similar than those from gadOn and parv. Given that parvalbumin-positive interneurons form a subset of Gad-positive interneurons (Pfeffer et al., 2013), this result is unexpected, and highlights how similar long-range input is to excitatory and inhibitory cell types in L2/3 V1.

Given this high similarity, we asked whether we could detect differences in input to cell types in L2/3 and to cells in other V1 layers. We therefore examined the difference between the presynaptic connectivity maps of the excitatory L2/3 gadOff brains and a class of excitatory L6 cells (ntsr1). Indeed, we found significantly larger GTD values between the gadOff and ntsr1 experiments (median = 0.030, IQR = 0.013, n = 12) (**Fig. 1d**, right), than either the pairs from gadOff and gadOn experiments (p = 1.3 x 10^−5^; rank-sum test) or the pairs from gadOn and parv experiments (p = 2.0 x 10^−6^), indicating input connectivity to excitatory and inhibitory cell types in L2/3 is highly homogeneous when compared to the input to cell classes in other layers.

We next sought to examine whether differences in input connectivity to different V1 cell types existed, using a finer-grained analytical approach. We analysed the same three groups as for the GTD analysis (gadOff vs. gadOn; gadOn vs. parv; and gadOff vs. ntsr1). Firstly, for each pair, the hierarchical segmentation ontology was simplified by excluding regions with no, or very few, presynaptic neurons (**Supplementary Fig. 3**; see Methods). Next, by traversing the resulting ontology from top to bottom, each brain region comprising at least two subregions in which cells were reliably detected was studied. Rather than perform multiple statistical comparisons on each of these brain regions, which may be susceptible to false-positives, we first applied a vector method to determine whether the overall distribution of inputs among its child subregions differed between target cell types (**Supplementary Fig. 4**; see Methods). If this difference was significant, each child subregion was then examined, using a t-test to determine whether the fraction of cells in the child subregion was different between the two target cell types. Importantly, at each step, the fraction of cells in the child subregion was expressed as a fraction of total cells in the parent region, rather than the overall total number of labelled cells across the brain, eliminating positive dependence in the hierarchy.

As suggested by the GTD analysis above, and despite experimental groups showing significant input from many brain regions (**Supplementary Fig. 5**), the overall input to the gadOff and gadOn groups was highly similar, with only two regions in the entire ontology showing significant differences in fractional input connectivity: visual cortex (VIS) and somatosensory cortex (SS) (**Fig. 2a,b**). Within the isocortex, excitatory L2/3 cells in the gadOff line received a greater fraction of input from the visual cortex (including primary and higher areas) than inhibitory gadOn cells (gadOff: 95.7 ± 1.3%; gadOn: 82.3 ± 1.6%; p = 0.001), and a lower fraction from SS (gadOff: 0.14 ± 0.07%; gadOn: 0.66 ± 0.15%; p = 0.04). However, no other significant differences were found in the hierarchy, either cortically or subcortically. Similarly, the two L2/3 inhibitory lines (gadOff and parv) showed very few differences, with the only regions differing significantly being in the subregions of the retrosplenial cortex (RSP). Namely, a greater fraction of RSP input to parvalbumin-positive VISp neurons came from RSPagl, whereas a larger fraction of input to the superset of neurons in the gadOn line came from RSPv.

**Figure 2.**
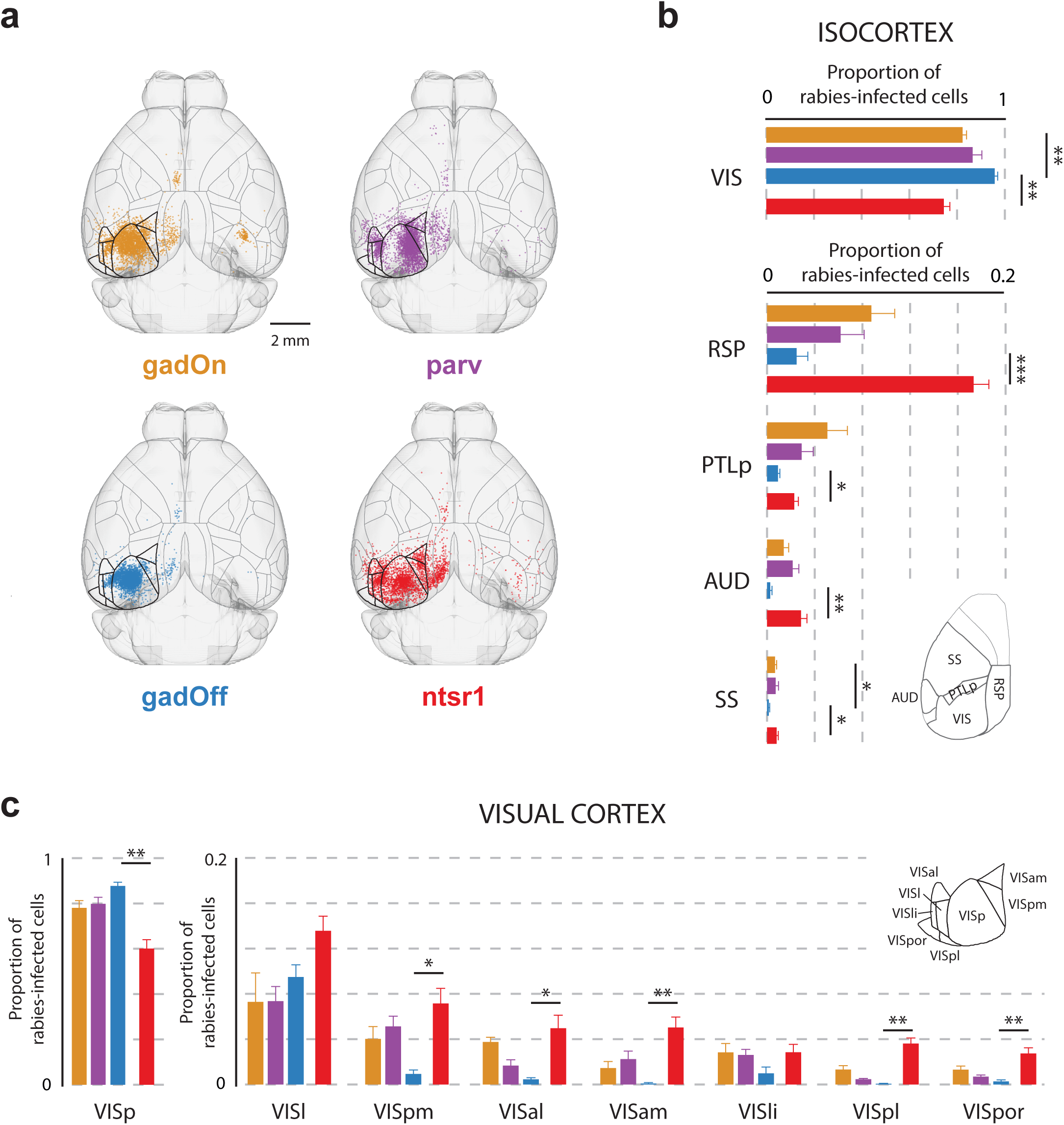
Distribution of presynaptic input from isocortex to primary visual cortex (VISp). (**a**) Horizontal projections of a template brain, showing the positions, after registration, of presynaptically connected cells within the isocortex, for each mouse line. The number of cells displayed for each line is the same, and has been normalized to the line with the fewest inputs (gadOff) by randomly sampling from all isocortex inputs. Both contra- and ipsilateral inputs are displayed, although only ipsilateral inputs were used for analysis. Horizontal surface projection of the segmentation is shown. Regions comprising visual cortex are highlighted with thicker outlines. (**b**) Proportion of labelled rabies-infected cells in ipsilateral top-level subregions of the isocortex. Inset, location of RSP, PTLp, AUD and SS on the horizonal surface projection of the segmentation. (**c**) Proportion of labelled cells in subregions of the visual cortex (VISp and higher visual areas). Inset, location of each visual area in the horizontal segmentation.

On the other hand, significant differences in fractional input to excitatory L2/3 cells and excitatory L6 ntsr1 cells were found in several regions across the brain (**Fig. 2,3**). From the isocortex, L6 ntsr1 cells received a significantly greater fraction of input from outside of visual cortex than L2/3 gadOff cells (fraction of isocortex presynaptic inputs in VIS; gadOff: 95.7 ± 1.3%; ntsr1: 74.5 ± 2.5%; p = 0.001, t-test), including a very large fraction from RSP (gadOff: 2.5 ± 0.9%; vs. ntsr1: 17.3 ± 1.3%; p = 0.0003) (**Fig. 2b**). Within visual cortical regions, again L6 ntsr1 cells received a lower fraction of local primary visual cortex (VISp) inputs than L2/3 gadOff cells (**Fig. 2c**) (gadOff: 87.7 ± 1.6% of VIS inputs from VISp; vs ntsr1: 60.0 ± 3.9%; p = 0.002). Indeed, in all secondary visual areas (VISpm, VISal, VISam, VISpl, VISpor) bar two (VISl and VISli) L6 received significantly more input than L2/3 gadOff cells (**Fig. 2c**). These data suggest that, although both L2/3 and L6 receive a wide range of inputs from similar cortical regions, L6 ntsr1 cells may integrate relatively more input from cortical regions outside VISp than superficial layers, which pools inputs more locally.

**Figure 3.**
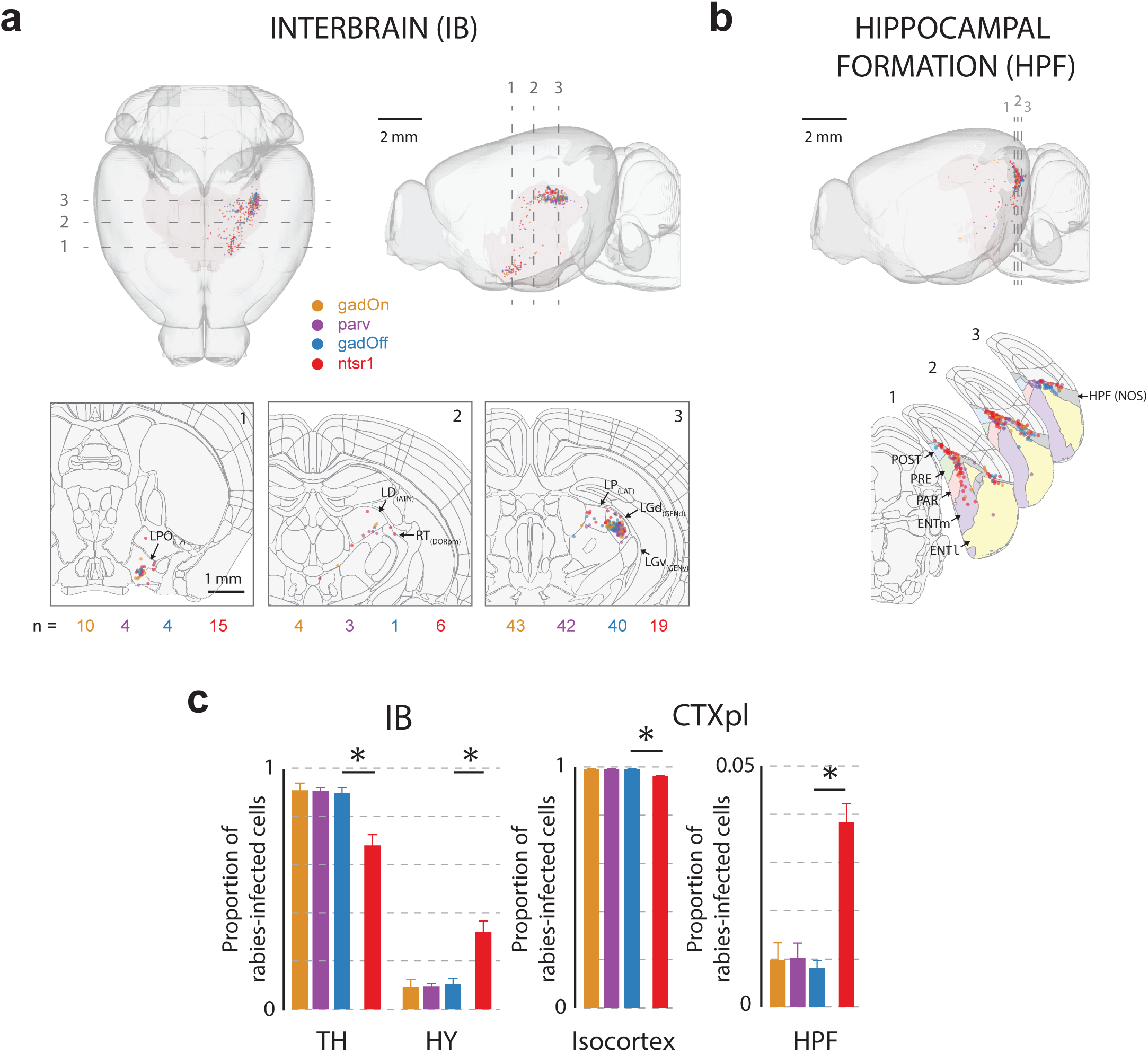
Distribution of presynaptic input from interbrain (IB) and hippocampal formation (HPF) to VISp. (**a**) Top, horizontal and sagittal projections, showing the distribution of labelled rabies-infected cells within the IB. Dashed lines indicate location of cross-section images, below. Bottom, coronal cross-sections of the template brain at three locations, with the position of labelled cells in the interbrain. Brain regions are outlined, and cells ±20µm anterior-posterior of the cross-section location are shown. Lateral preoptic area (LPO), lateral dorsal nucleus (LD), reticular nucleus (RT), lateral posterior nucleus (LP), and dorsal (LGd) parts of the lateral geniculate complex are indicated (in brackets, the parent regions of these regions). (**b**) Sagittal projection showing the distribution of labelled cells within the HPF. Dashed lines indicate location of cross-section images, below. Below, coronal cross-sections of the template brain at three locations, with the position of labelled HPF cells and brain regions outlined (includes cells ±10µm anterior-posterior of the cross-section location). Pre- (PRE), post- (POST) and para- (PARA) subiculum, as well as lateral (ENTl) and medial (ENTm) entorhinal cortex are indicated. The dark grey region indicates a region labelled HPF with no further subdivision (NOS – not otherwise specified). (**c**) Proportion of rabies-infected cells within the subregions of the interbrain (IB) and cortical plate (CTXpl).

Outside of the isocortex, although no significant differences were seen in the fractional input to different types of L2/3 cell, several regions differed significantly in the input they provided to L6 ntsr1 versus L2/3 gadOff (**Fig. 3**). For instance, interbrain (IB) (**Fig. 3a,c**) inputs on to L2/3 gadOff cells were more likely to originate from thalamus (gadOff: 89.5 ± 2.3% vs. ntsr1: 67.9 ± 4.5%; p = 0.01), and, accordingly, L6 cells received significantly higher fraction of their IB input from regions of the hypothalamus (**Fig. 3c**). L6 ntsr1 cells also received a greater fraction of cortical plate inputs from the hippocampal formation (HPF) than L2/3 (gadOff: 0.81 ± 0.16% vs. ntsr1: 3.83 ± 0.39%; p = 0.002; **Fig. 3b,c**) including inputs from CA1 and the post-, pre- and parasubiculum, as well as lateral and medial entorhinal cortex.

## Discussion

Several recent studies in the neocortex and midbrain have examined the brain-wide distribution of direct inputs onto different cell classes (visual cortex: Vélez-Fort et al., 2014; Kim et al., 2015; Leinweber et al., 2017; Liang et al., 2019; prefrontal cortex: Ährlund-Richter et al., 2019; Sun et al., 2019; somatosensory cortex: DeNardo et al., 2015; Wall et al., 2016; Hafner et al., 2019; Zolnik et al., 2020; ventral tegmental area: Beier et al., 2015, 2019). Many of these studies have shown only modest effects of cell class on long-range connectivity, with all cell classes in a region generally receiving projections from similar distant regions. Here we have examined the distribution of whole-brain, monosynaptic input to excitatory and inhibitory neurons in L2/3 of V1 using rabies-viral tracing. Consistent with these previous studies it appears inhibitory and excitatory neurons in L2/3 of mouse V1 receive highly overlapping inputs from distal brain areas.

However, in contrast to previous studies (e.g. Ährlund-Richter et al., 2019; Wall et al., 2016) rather than using multiple Cre-lines to distinguish subpopulations of, for example, GABAergic cells, we have rather studied excitatory and inhibitory classes more generally, using Cre-Off and Cre-On AAVs in Gad2-Cre mice. Thus, this study has not asked whether subtypes of inhibitory or excitatory cells might be differentially innervated, but whether the two classes receive anatomically segregated distant inputs. For comparison, we also examined input to parvalbumin-cells, the largest known subtype of GABAergic neuron in the neocortex (Markram et al., 2004). Although the differences were small, there were noticeable qualitative trends in the fraction of inputs to the different classes, suggesting the potential for more subtle diversity in the inhibitory population than our study could detect.

It is clear from these data that deep layer ntsr1 cells are readily distinguishable from upper layer cells based on the profile of their long-range input. On a global scale, and although we restricted the majority of our analysis to ipsilateral input, ntsr1 cells received a significantly higher fraction of contralateral inputs than gadOff and parv cells, whereas none of the L2/3 lines were significantly different to each other (**Supplementary Fig. 1**). Further, the anatomical weighting of RSP input, which carry head-motion related signals to deep layers (Vélez-Fort et al., 2018) is substantially higher than to upper layers (Leinweber et al., 2017; Vélez-Fort et al., 2014; Chaplin and Margrie, 2020). Indeed, L6 received relatively large amounts of direct input from regions in the hippocampal formation, such as CA1 (Cenquizca and Swanson, 2007) and presubiculum, parasubiculum and postsubiculum (Vogt and Miller, 1983), which contain head direction responses (Taube et al., 1990), and which are thought to strongly contribute to grid cell representations in entorhinal cortex (Moser et al., 2008). Therefore, it appears processing in upper and deeper layers utilizes information from functionally distinct long-range inputs.

Although these data indicate that both excitatory and inhibitory neurons in L2/3 receive similar information, it is important to recognize that physiological processing of that information will depend on several other key factors, such as the relative weight and dynamics of these long-range synaptic connections. Also, it will be important to determine whether individual inhibitory and excitatory cells share input from the same presynaptic cells, or rather process information in parallel, from distinct presynaptic populations.

Despite similar regions connecting in similar proportions to cell types in L2/3, different cell classes within localized regions of V1 possess distinct visual and behaviourally-relevant response properties (Kerlin et al., 2010; Hofer et al., 2011; Khan et al., 2018). Thus, major differences in the response profiles of these cell classes likely arise from a combination of diversity in synaptic physiology (Reyes et al., 1998), intrinsic biophysical properties of the target cells (Brown et al., 2019) and local circuit connectivity (Bock et al., 2011; Ko et al., 2011; Packer and Yuste, 2011) rather than long-range, inter-areal connectivity.

In summary, our data suggest that L2/3 excitatory and inhibitory populations in visual cortex receive strongly overlapping long-range inputs. These inputs are relatively local when compared to the input to ntsr1 cells of L6, which integrate more non-visual, multi-modal input. Thus, in upper layers within the cortical column, processing of similar long-range information in upper layers likely combines with translaminar processing to shape diverse cortical function (D’Souza and Burkhalter, 2017; Adesnik and Naka, 2018).

## Acknowledgements

We are grateful to the support staff of the Neurobiological Research Facility and thank Larry Swanson and Mateo Vélez-Fort for providing comments on earlier drafts of this manuscript. This work was funded by The Gatsby Charitable Foundation and The Wellcome Trust (214333/Z/18/Z) (T.W.M.).

## Contributions

A.P.Y.B. and T.W.M. designed the experiments. A.P.Y.B. performed the experiments. A.P.Y.B. and L.C. analysed the data. A.P.Y.B., L.C. and T.W.M. wrote the manuscript.

---

**Supplementary Figure 1.**
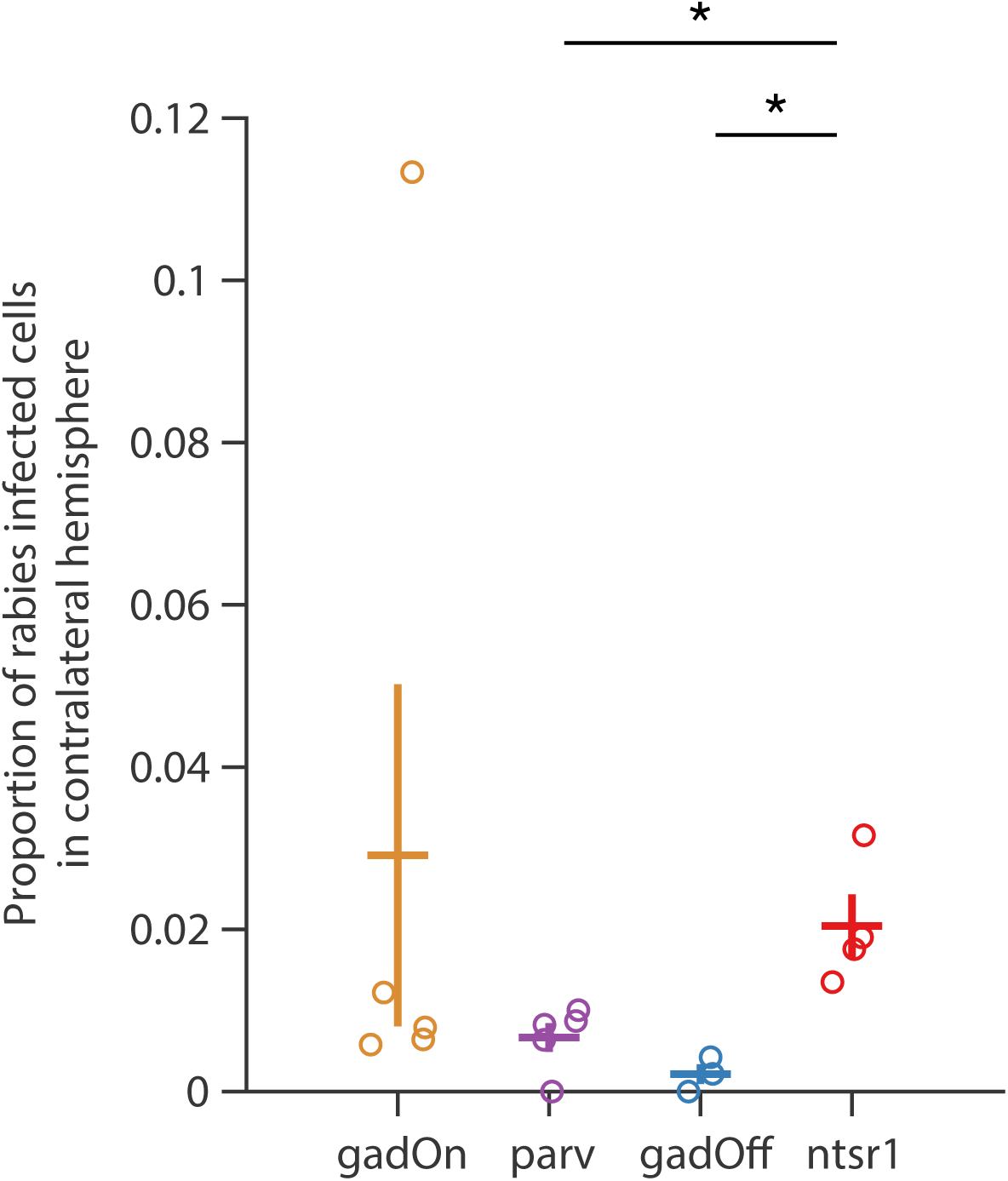
Fraction of cells in the contralateral hemisphere. Proportion of the total number of rabies-infected cells in whole brain, found in the contralateral hemisphere, for each mouse line. The proportion for each individual brain is displayed as a dot, and the mean and SEM for each mouse line are shown. The ntsr1 line had significantly more cells in contralateral hemisphere than the parv (p = 0.011) or gadOff lines (p = 0.012; unpaired t-tests).

**Supplementary Figure 2.**
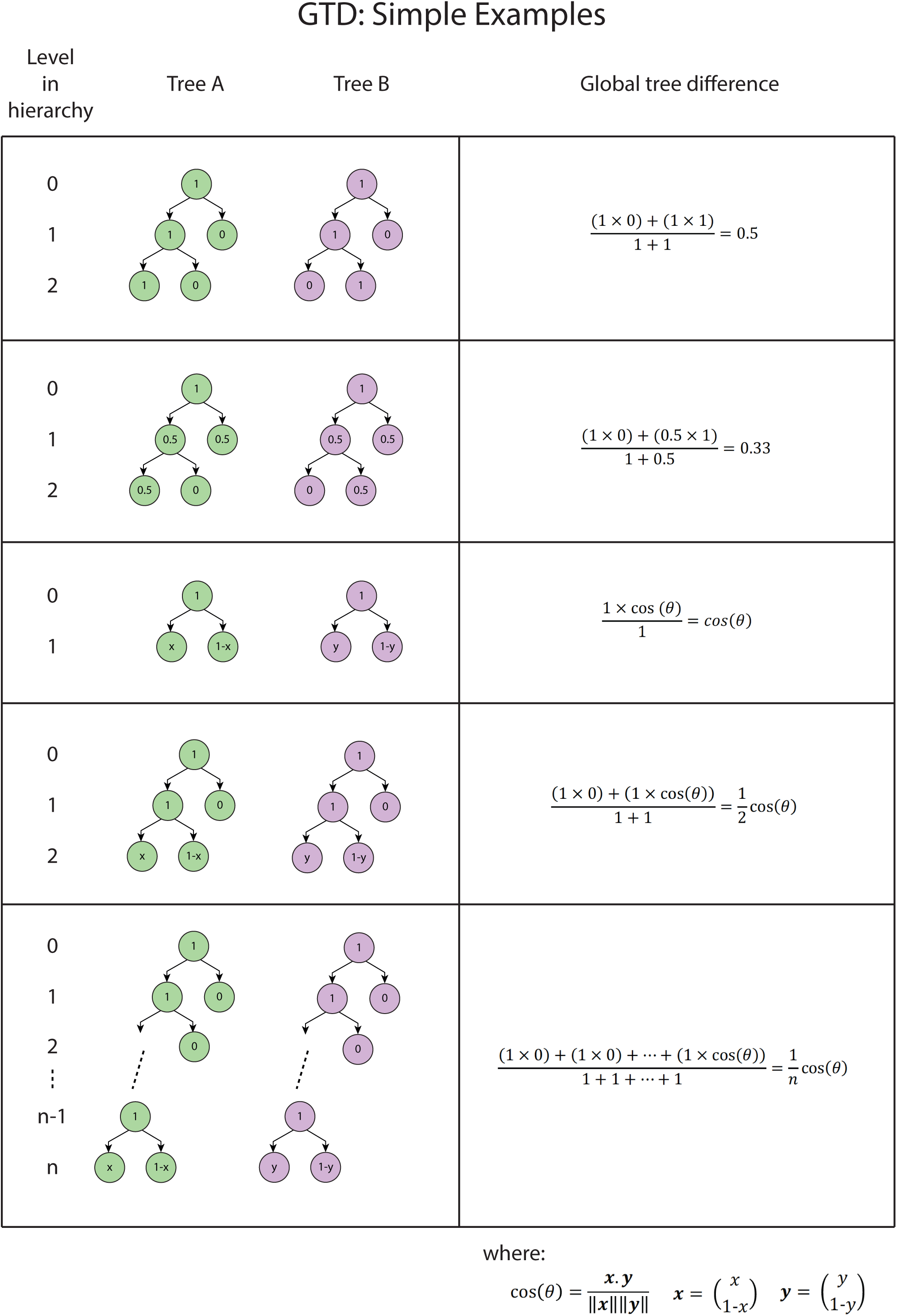
Examples of simple trees and their global tree difference. Several examples of simple tree structure and their global tree difference values.

**Supplementary Figure 3.**
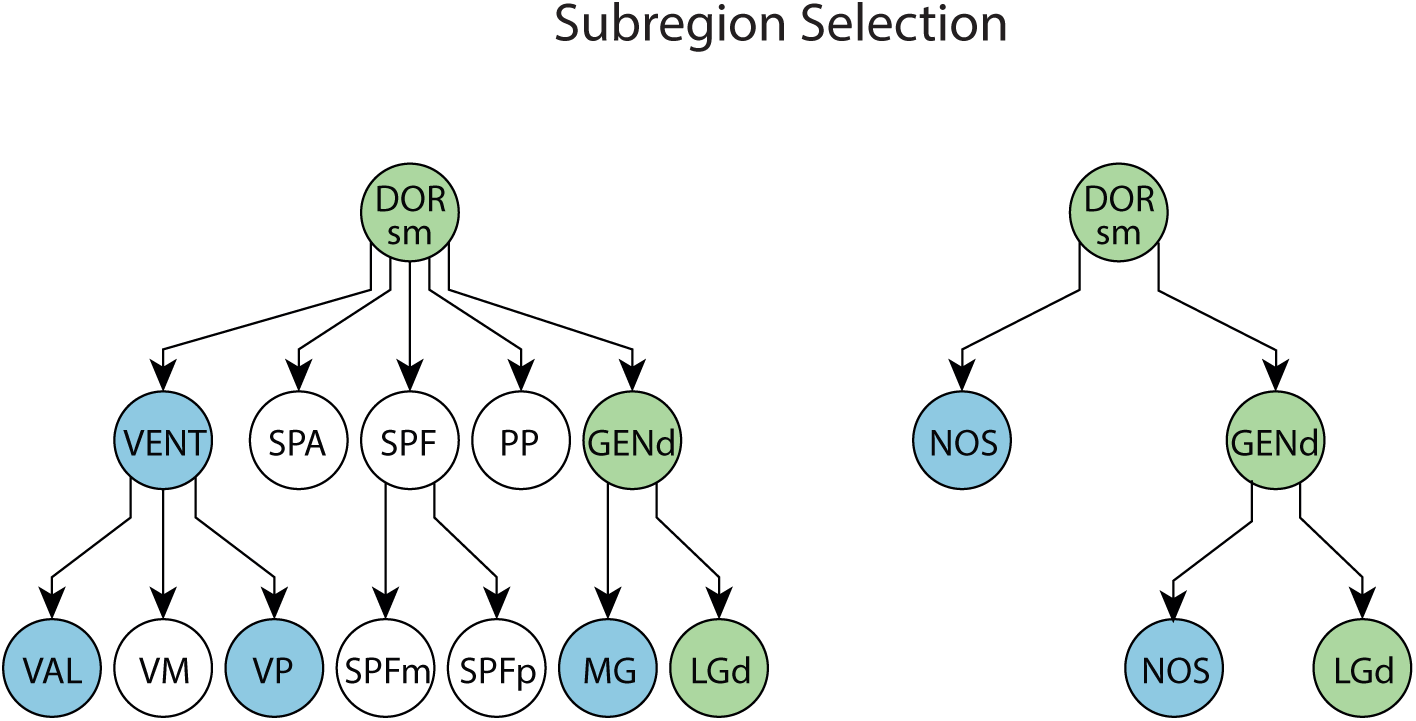
Selection of regions for analysis. An example of applying the subregion selection policy to the dataset. Left, a portion of the full ontology is shown from DORsm in the thalamus. Regions which meet the selection criteria (see Methods) are coloured in green. Regions containing some neurons but not meeting criteria for inclusion are shown in blue; regions with no neurons are shown as empty circles. Right, amalgamated ontology. Subregions not meeting criteria (blue) have been combined into a ‘not otherwise specified’ (NOS) subregion (which also included cells segmented to the parent but not any of its subregions). NOS subregions are included in the vector analysis stage, but are not further analysed.

**Supplementary Figure 4.**
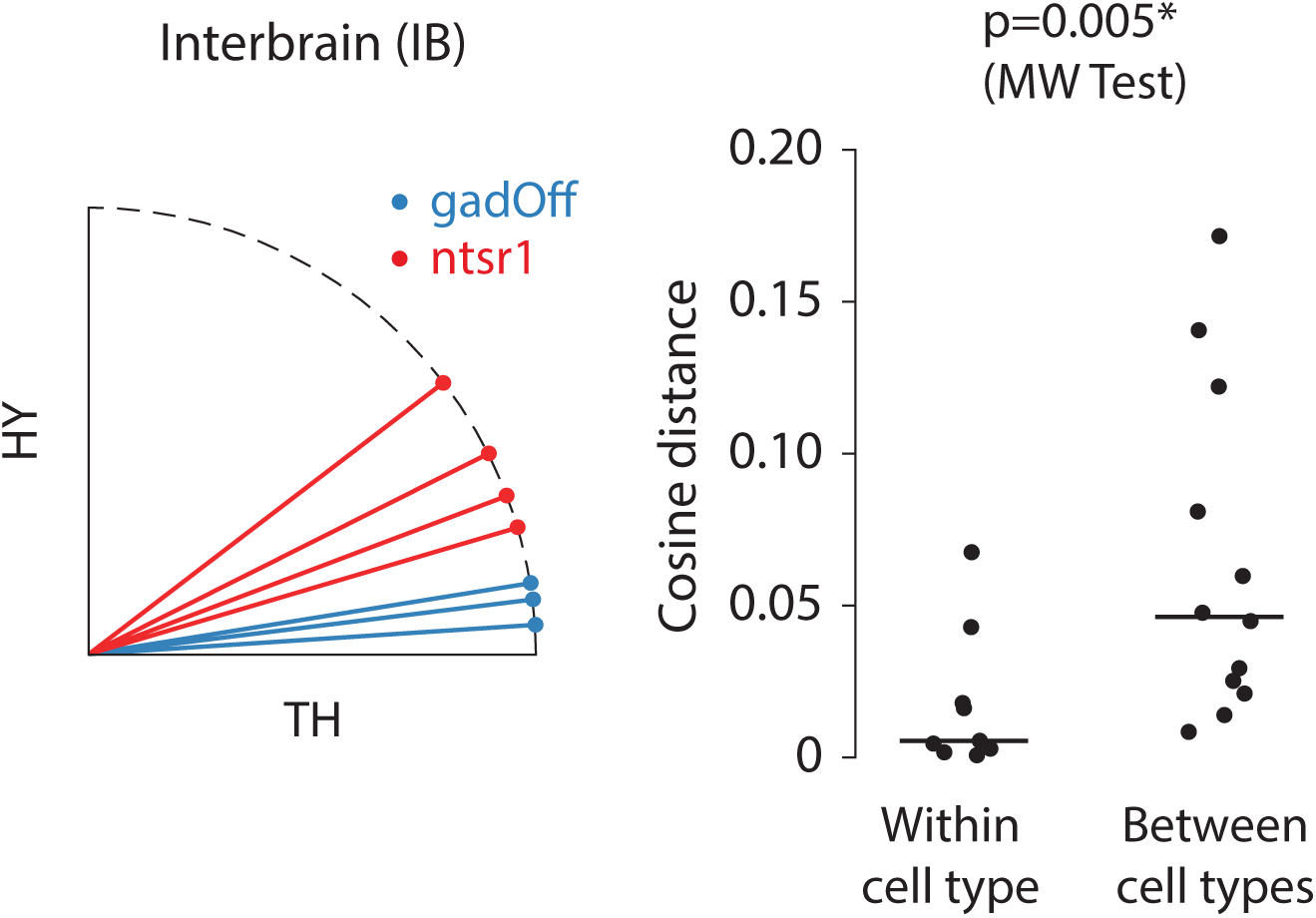
Vector-space subregion analysis. An example of the vector-space subregion analysis applied to the subregions of interbrain (composed of thalamus (TH) and hypothalamus (HY)). Left, the fractional composition of subregions for each experiment is plotted as a vector of unit length. Vectors are shown for the gadOff and ntsr1 groups. NOS fractions have not been shown, although they are always used in the analysis where present. Right, comparison of cosine distance between all pairs of vectors within mouse lines and between mouse lines.

**Supplementary Figure 5.**
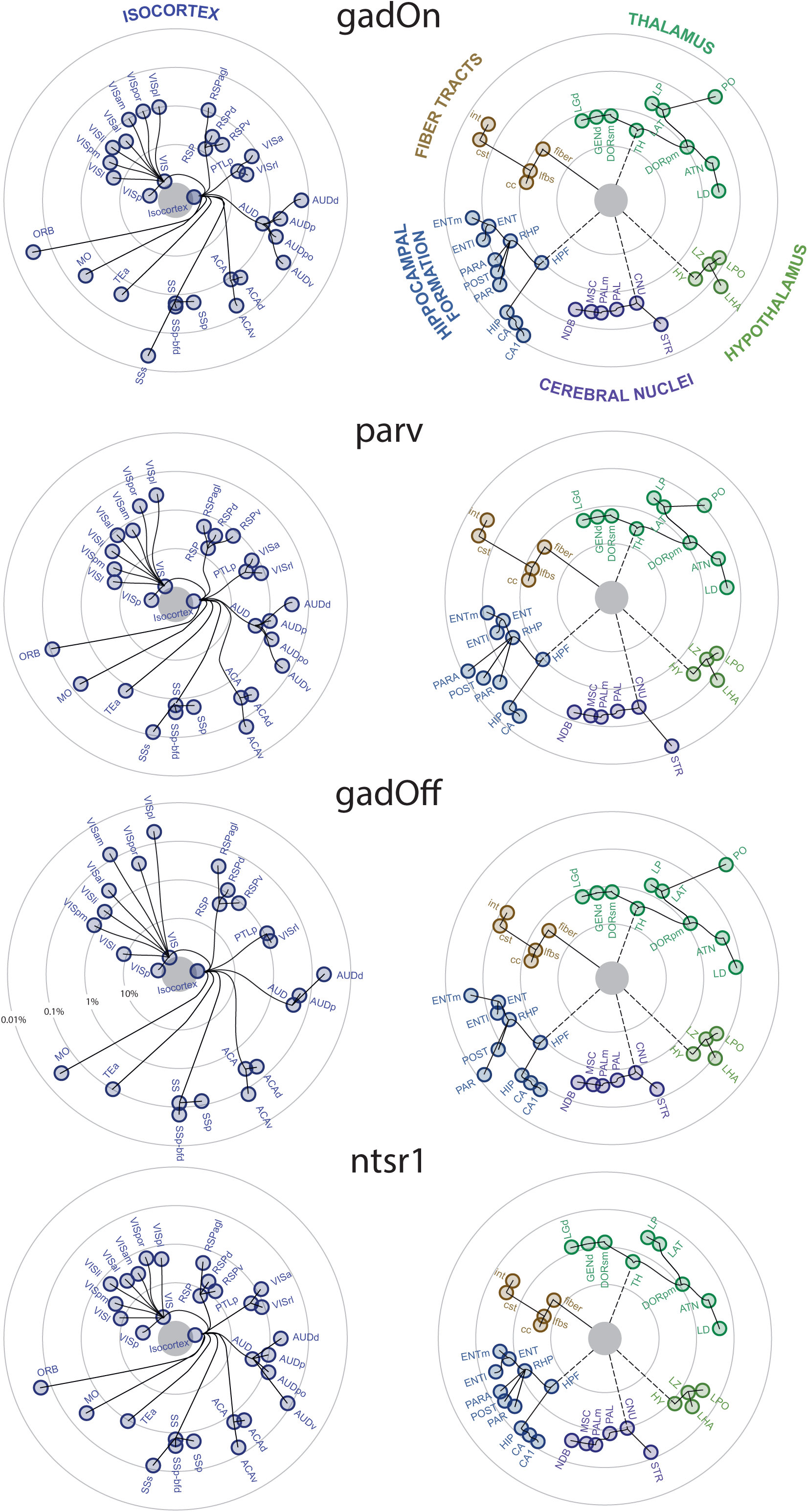
Topography of presynaptic input to cell types of primary visual cortex. Radial dendrograms showing the topography of average presynaptic connectivity to V1 for gadOn, parv, gadOff and ntsr1. Each node on the circular plot indicates a brain region, and the distance of the node from the edge of the centre circle indicates the average fraction of total rabies-infected cells found in that region. Lines connect brain regions appearing in the same branch of the Allen Reference Atlas ontology. Labels are consistent with the Allen Reference Atlas nomenclature.

## Methods

### Transgenic Animals

Conventional AAV-assisted rabies virus tracing was carried out in Gad2-Cre (interneurons; Jax 10802), Parv-Cre (parvalbumin positive interneurons; Jax 008069) and Ntsr1-Cre (layer 6 cortico-thalamic cells; GenSAT 030648-UCD) mice. Ntsr1 data were taken from (Vélez-Fort et al., 2014), but cell counts were reanalysed in the same manner as the other datasets. To target layer 2/3 excitatory and inhibitory neurons, we used superficial injections of Cre-inactivated or Cre-activated viral constructs in Gad2-Cre animals (here termed “gadOff’” and “gadOn”, respectively). Both male and female adult animals (P35-56) were used. All animals were bred on to a C57/BL6J background.

### Surgical Procedures

All experiments were performed in accordance with the UK Home Office regulations (Animal (Scientific Procedures) Act 1986) and the Animal Welfare and Ethical Review Body (AWERB). Surgical procedures were carried out under anaesthesia, using a mix of fentanyl (0.05mg/kg) / midazolam (5mg/kg) / medetomidine (0.5mg/kg). Anaesthesia was reversed at the end of the procedure using naloxone (1.2mg/kg) / flumazenil (0.5mg/kg) / atipamezole (2.5mg/kg) where necessary. During all procedures, both eyes were protected by the application of ointment (Maxitrol, Alcon).

The scalp overlying the left primary visual cortex was shaved, cleaned, and the animal was placed on a warming mat controlled by an internal temperature probe to ensure core body temperature did not fall below 36°C. After incising the scalp, a small (approx. 0.5mm diameter) craniotomy was then drilled over the monocular portion of V1 at 0.5mm rostral and 2.3mm lateral to lambda using a dental drill (Osada Electric, Japan) with a 300µm burr (Cookson). Viral injection was then performed (see below), following which the craniotomy was filled with a silicone elastomer (Kwik-Cast, WPI), and the scalp overlying closed using either sutures or cyanoacrylate glue. The animal was then recovered, using a naloxone / flumazenil / atipamezole mix, and replaced in the home cage.

Subsequent viral injections were performed in the same way, with the exception of the craniotomy; the silicon plug was quite easy to remove and there was little connective tissue regrowth between surgeries.

### Virus Injection

Viral injections were carried out using a borosilicate pipette with a long, narrow shank, with a tip broken to a diameter of around 15-20µm. In some cases, a motorised injection system (Nanoject, Drummond) was used, however most injections were carried out using a 2ml air-filled syringe for pressure, whilst visualising the meniscus of the viral suspension under high power magnification.

Injections in Ntsr1-Cre mice were carried out at a depth corresponding to the middle of L6 (∼690µm below pia). In the gadOff, gadOn, and PV-Cre mice (targeting layer 2/3 neurons) injections were performed close to the border of layers 1 and 2 (around 120µm below the pia), to minimise the transfection of deeper neurons. For conventional (Cre-On) tracing, a mix of pAAV hSyn-Flex RG-cerulean (Addgene 98221) and pAAV-EF1a-FLEX-GT (gift from Edward Callaway; Addgene 26198) was used in a 2:1 ratio. For Cre-Off tracing, a mix of pAAV-EF1a cre off Cer-E2A-RG (Addgene 126471) and pAAV-EF1a-cre off EGFP-2A-TVA (Addgene 126470) was used. The AAV mix appropriate for the particular experiment was injected over 1-2 minutes in short pulses. Approximately 5-20nL was injected for each experiment. Following the injection, the pattern of blood vessels was imaged allowing for easy identification of the injection site.

Two to three days later, rabies virus (RV, 100-150nL) was injected at the same site, using blood vessel patterns at landmarks. Smaller volumes of AAV as compared to the RV were used, in order to cause primary infection of as many transfected neurons as possible, whilst at the same time restricting the number of host neurons so as to make cell counting practicable.

### Perfusion

Animals were perfused 10-14 days after rabies infection. Animals were deeply anaesthetized using a ketamine (200mg/kg) / xylazine (20mg/kg) mixture, and a blunt needle placed in the left ventricle, whilst an incision was made in the right atrium. Blood was first cleared using 100mM phosphate buffered saline (PBS); once clear, the animal was perfused with saline containing 4% paraformaldehyde. Once fully fixed, the head was removed and the brain carefully dissected out. The brain was further fixed in 4% PFA overnight at 4°C, and then stored in 100mM PBS with 0.1% azide to prevent bacterial contamination, until ready for imaging.

### Imaging

On the day of imaging, brains were removed from the PBS and carefully dried. They were embedded in agarose (4%) using a custom alignment mould to ensure that brain was perpendicular to the imaging axis.

After trimming, the agarose blocks were transferred to the serial two photon microscope containing an integrated vibrating microtome and motorized x-y-z stage (Osten and Margrie, 2013; Ragan et al., 2012). Two-photon imaging was carried out using 930nm or 800nm illumination. Images were acquired with a 1μm pixel size, and 5μm plane spacing. 10 optical planes were acquired over a depth of 45μm in total. In order to image the entire brain, mosaic scanning was employed, in a 13×9 grid with each tile measuring 1080 pixels square, with 5% overlap.

After imaging each mosaic tile at all 10 optical planes, the sample was automatically transferred to a microtome which removed a 50μm slice, allowing for imaging of the subsequent portions of the sample, resulting in full 3D imaging with a voxel size of 1μm in-plane, and 5μm axially.

### Data analysis

#### Segmentation and cell counting

Individual tiles were preprocessed to correct for uneven illumination, and then stitched into 2D whole-brain coronal slices using custom routines written in Python. Slices were then loaded as 3D stacks and inspected using the MaSIV package (available at https://github.com/SainsburyWellcomeCentre/masiv) designed to load a downsampled image stack fully in to memory, whilst dynamically loading full resolution images from disk as needed. Cells were manually counted in each brain using a cell counter plugin, which reports the precise position of each cell. To aid counting in cases where perfusion was imperfect, images underwent linear unmixing using emission spectra derived from pure samples of GFP, mCherry, and blood.

Rabies-infected cells were identified manually as smooth objects with diameter of 8-30um and fluorescence only in the red channel (or red and green channels, in the case of host neurons in the primary injection site). Neuronal processes were often visible, further aiding identification.

Image stacks were downsampled and aligned to the Allen Reference Atlas using the aMAP algorithm described by Niedworok et al. (2016) (available at https://github.com/SainsburyWellcomeCentre/aMAP), which is based upon NiftyReg (Modat et al., 2010). Cell positions were then projected on to the reference space, and segmented automatically.

#### Simplification of the ontology

Presynaptic neurons were segmented according to the Allen Reference Atlas (2015) ontology. We applied the following selection criteria of regions for further analysis. Areas containing no neurons were excluded from analysis. Additionally, areas were only included for analysis provided the following conditions were met, in at least one line: first, the area must have at least 5 neurons, on average, across all brains in one line; second, the area must have at least 1 neuron in the majority (>50%) brains in one or more lines. In order to analyse and compare how presynaptic cells were distributed between the subregions of an area, we separated the area into those subregions which passed the above criteria and a “not otherwise specified” (NOS) subregion. The NOS subregion was the amalgamation of all the cells in the subregions not passing the above criteria, as well as cells which were assigned to the parent area directly (i.e. not assigned to a particular subregion). The NOS subregion was not analysed for differences between lines (t-tests below), but did contribute when computing the vector space subregion analysis (see below). This step, as well as the vector space subregion analysis and subsequent t-tests, were performed separately on three groups: inter-laminar excitatory (gadOff and Ntsr1), inter-cell-class L2/3 (gadOff and gadOn) and inhibitory L2/3 (gadOn and Parv).

#### Vector space subregion analysis

Once simplified, each area with at least 2 remaining subregions (not including the NOS subregion) was first analysed in order to determine whether the fraction of cells in each subregion differed systematically by host line. This step reduces the number of subsequent t-tests required (see below).

For a given area containing *n* subregions, the fraction of cells in all subregions was expressed as an n-dimensional vector for each brain. The direction of this vector provides a complete description of the cellular subregional composition for an area in a single brain. The angle between any two such vectors defines a measure of the overall similarity of the vector directions. For data such as these, which are bounded in [0, 1] in all dimensions, the cosine of the angle between the two vectors is commonly used, varying from 0 for orthogonal vectors to 1 for vectors with identical direction.

Cosine distance was therefore calculated for all pairs of experiments from within the same line, and for all pairs of experiments from different lines. These two sets of distance measures were then compared using a one-tailed Mann-Whitney test. A significant result indicates that pairs of experiments within a line are more similar in their subregional composition than pairs of experiments between lines.

#### Difference by line of individual subregion fractions

In regions in which the vector space subregion analysis was significant (p < 0.05), we next examined each child subregion of that parent region (not including the NOS region), by using an unpaired t-test to compare the fraction of cells in each line.

#### Global Tree Difference

The Global Tree Difference was defined to be a scalar value by which a pair of segmentations can be compared (see Fig. 1c and Supplementary Fig. 2 for examples). It is defined as the average cosine distance of subregional composition for each parent region, weighted by the fractional input in the region, and normalized to the total number of cells in the two brains.

For each region in the ontology, the cosine distance between the two experiments is first calculated as outlined above. This is then weighted by the average of the fractional cell counts (the number of labelled cells in the region and all subregions, divided by the total number of labelled cells in the entire hemisphere). Lastly, the metric is normalized by dividing by the total mean fractional connectivity, to give a value bounded in [0,1]. High values of GTD are hard to achieve, requiring orthogonal subregional composition at the highest levels of the hierarchy; therefore, in experiments in which differences are not gross, it is expected that GTD values will cluster near to 0.

## Notes

### Competing Interest Statement

The authors have declared no competing interest.

